# Widespread loss of sleep in independently evolved populations of wild-caught cavefish

**DOI:** 10.1101/2024.10.29.619241

**Authors:** Owen North, Lourdes Citlalli Maza-Castañeda, Aubrey E. Manning, Aakriti Rastogi, Andrew Gluesenkamp, Leah Gluesenkamp, Nathan Swanson, Jorge Hernández-Lozano, Marco A. Garduño-Sánchez, Jesús Alberto Díaz-Cruz, Johanna E. Kowalko, Suzanne E. McGaugh, Claudia Patricia Ornelas-García, Alex C. Keene

**Author notes:** Denotes equal contribution (Owen North, Lourdes Citlalli Maza-Castañeda, Aubrey E. Manning, Claudia Patricia Ornelas-García).

## Abstract

Sleep is an evolutionarily ancient and nearly universal behavior throughout the animal kingdom. Multiple cave-dwelling populations of the Mexican tetra, *Astyanax mexicanus*, have converged on sleep loss compared to river-dwelling surface fish. However, sleep has not been assessed in the vast majority of the 34 known *A. mexicanus* cave populations. Moreover, whether cavefish sleep less than surface individuals in their natural habitats is currently unknown. Analyzing the distribution of sleep loss and its relationship with other regressive traits in a phylogenetic framework is critical to inform the selective pressures across the different lineages. We measured sleep and locomotor activity in 15 distinct populations of *A. mexicanus,* including lineages that are broadly representative of the 34 cavefish populations identified to date. Sleep was drastically reduced in all cave and hybrid populations that were tested. A subset of caves contain hybrid populations of *A. mexicanus*, which show a broad range of eye and pigmentation phenotypes, yet have evolved near-complete loss of sleep. Mapping behavioral changes onto the phylogeny of *A. mexicanus* populations revealed that loss of sleep and elevated locomotor activity have evolved at least three times. Analysis of sleep in the wild confirms that the sleep loss phenotype observed in lab-reared fish is also present in the natural environment. Together, these findings reveal deep evolutionary convergence on sleep loss in cavefish and provide evidence for sleep loss as a primary trait contributing to cave adaptation.

## Introduction

Sleep is an ancient and nearly universal behavior throughout the animal kingdom (Allada and Siegel, 2008; Anafi et al., 2019; Campbell and Tobler, 1984; Keene and Duboue, 2018). While sleep duration and timing vary across species, sleep function appears to be highly conserved, with loss of sleep disrupting brain function and physiological homeostasis (Allada and Siegel, 2008; Arble et al., 2015). Despite the critical role of sleep in maintaining core biological functions, numerous species can forgo, or significantly alter, sleep for extended periods under ecologically advantageous conditions, such as extended foraging periods or mating seasons (Rattenborg and Ungurean, 2023; Siegel, 2009). Identifying the evolutionary and ecological factors influencing sleep duration between and within species is critical for understanding the function of sleep.

The Mexican cavefish, *Astyanax mexicanus*, is a model for investigating behavioral evolution (Keene and Appelbaum, 2019). *A. mexicanus* exists as eyed surface-dwelling populations and at least 34 known blind cave-dwelling populations (Cobham and Rohner, 2024; McGaugh et al., 2020). Within the past 200,000 to ∼5 million years, multiple colonizations of caves by eyed surface ancestors have resulted in multiple cave populations that are geographically and hydrologically isolated from each other (Casane and Rétaux, 2016; Gross, 2012; McGaugh et al., 2020; Swaminathan et al., 2024). Independently evolved cave populations have converged on numerous traits including eye degeneration, albinism, hyperphagia, and sleep loss (McGaugh et al., 2020; Swaminathan et al., 2024). To date, sleep loss in cavefish has been documented to have evolved multiple times, suggesting that shared evolutionary pressures between cave populations contribute to convergence on sleep loss (Duboué et al., 2011; Moran et al., 2022b; Yoshizawa et al., 2015). Surface and cave populations are interfertile, and this has been exploited in laboratory settings to study trait segregation in surface-cave hybrid fish, revealing that distinct genetic architecture regulates morphological changes and sleep (Casane and Rétaux, 2016). These lab-based studies have revealed that evolved genetic and neural changes associated with sleep loss are also shared with human pathologies of sleep (Jaggard et al., 2018; Mack et al., 2021).

A unique strength of the cavefish system is the ability to study naturally occurring genetic variation across 34 cavefish populations (Miranda-Gamboa et al., 2023). While laboratory-based studies have largely been restricted to three populations of cavefish, genomic analysis of wild-caught cave populations has provided a detailed phylogeny and genomic markers of selection that are shared across distinct cavefish lineages (Bradic et al., 2013; Moran et al., 2023; Ornelas-García et al., 2008). Genes associated with sleep and circadian rhythms are enriched for markers of selection across independent phylogenies of cavefish, raising the possibility that changes in these processes are widespread, and that loss of sleep may be critical for cave adaptation (Moran et al., 2022a). However, no study to date that has analyzed sleep loss across a known phylogeny.

Genomic analysis has shown that several cave populations have experienced introgression with surface fish, allowing the 34 cave populations to be broadly classified as either non-introgressed or hybrid at the genomic level (Garduño-Sánchez et al., 2023; Moran et al., 2023). The presence of naturally-occurring hybrids provides a powerful opportunity to study the evolution of complex traits. Previous studies in a single hybrid population, Chica cave, have shown variation in sleep, eye size, and pigmentation, and evidence of convergent evolution in a circadian clock gene across multiple subterranean species (Moran et al., 2022b). Therefore, environmental heterogeneity (i.e., variation among pools within this population) can affect adaptive processes in cave environments (Moran et al., 2023).

Limited accessibility to many cave populations for behavioral analysis has significantly hindered progress in this field. The challenges associated with obtaining living specimens that can be studied in a laboratory have resulted in most behavioral analyses focusing on a small number of populations (Bibliowicz et al., 2013; Espinasa et al., 2021; McGaugh et al., 2020). Furthermore, while a small number of studies have examined behavior in the cave setting, sleep analysis requires multi-day recordings, and therefore, sleep has not been studied in wild *A. mexicanus* or any other fish species to date (Keene and Appelbaum, 2019). Examining the evolution of sleep across *A. mexicanus* populations provides a unique opportunity to reveal ecological factors that shape sleep regulation in the Mexican tetra.

Here, we investigate sleep and activity patterns across 15 wild-caught cave and surface populations, nearly all of which have not been studied for sleep or behavioral analysis. These studies reveal widespread loss of sleep across cavefish populations and show that this trait has independently emerged at least three times during evolution. The sleep loss identified under standard laboratory conditions is replicated in the wild, supporting the notion that sleep loss is adaptive for cave-dwelling *A. mexicanus*.

## Results

To determine whether sleep loss identified in laboratory-raised cave populations is a consistent feature of cave populations, we collected wild-caught *A. mexicanus* from independent lineages. All fish were wild-caught as adults and maintained in the laboratory prior to behavioral analysis (Supplemental Table 1). The populations of wild-caught *A. mexicanus* included fish from four surface populations (Bocatoma, Cueva del Rancho Viejo Spring, Micos River, and Rascón), five hybrid cave populations (Arroyo, Caballo Moro, Chiquitita, Cave Río Subterraneo, and Toro), and six cave populations (Escondido, Japonés, Pachón, Pichijumo, Piedras, and Tigre) (Fig 1A, B)(Garduño-Sánchez et al., 2023; Moran et al., 2023). Phylogenomic reconstructions validated that cave and hybrid populations derive from at least two distinct lineages, and at least three independent clades (Garduño-Sánchez et al., 2023; Kellermeyer et al., 2024; Moran et al., 2022b), that are representative of all 34 cavefish populations known to date (Fig 1). These fish are phenotypically diverse with clearly visible eye regression and loss of pigmentation observed in individuals from hybrid and cave populations (Fig 1B), providing the opportunity to measure evolved changes in sleep across Mexican tetra lineages.

**Figure 1.**
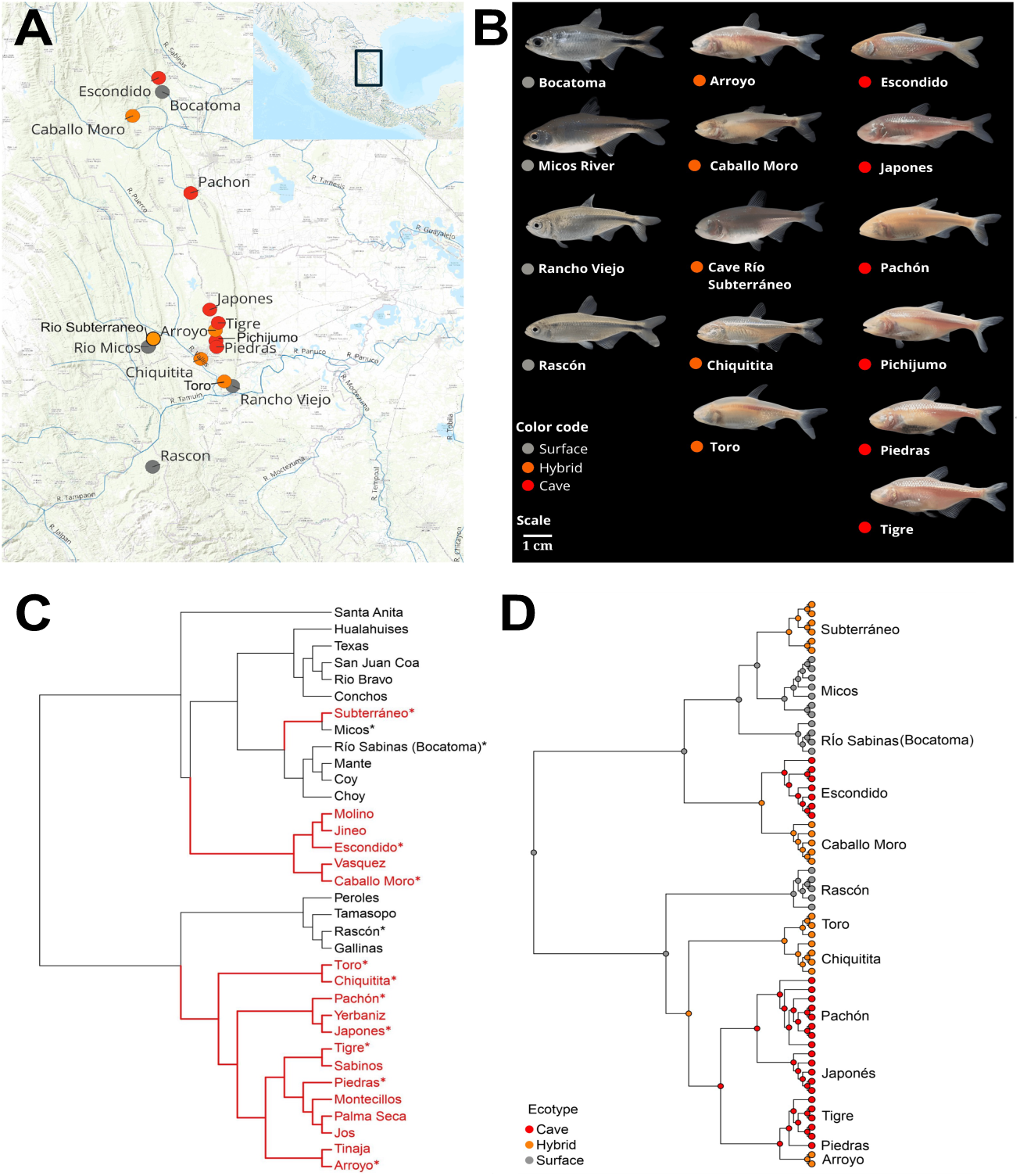
A) Map with the 15 populations used in the present study. **B)** Images depicting an individual fish from each independent population tested. Each population was characterized by ecotype: surface (gray), hybrid (orange), and cave (red). **C)** Complete phylogeny of *Astyanax mexicanus,* highlighting cave populations in red based on Garduño-Sánchez et al. 2023. Terminals marked with an asterisk, correspond to populations included in this study. **D)** Phylogenetic reconstruction based on a Maximum likelihood reconstruction of the cave and surface populations of A. mexicanus, based on 150,260 SNPs. The topology was edited to include the 62 terminals to visualize the sleep results obtained in the study. Populations were categorized as surface (gray), hybrid (orange), or cave (red). The three groups were defined based on the evidence of admixture observed in both Garduño-Sánchez et al. 2024, and Moran et al. 2024.

To quantify sleep, we recorded activity of individually housed fish over 24 hours. We then analyzed sleep and waking activity across the 15 distinct wild-caught populations. Wild-caught fish from the four surface populations slept on average 3-6 hours per day, roughly comparable to previous analyses of sleep in lab-reared adult surface fish (Jaggard et al., 2018; Yoshizawa et al., 2015). Conversely, sleep was greatly reduced or absent in nearly all fish from all hybrid and cave populations tested (Fig. 2A). The five hybrid and six cavefish populations slept significantly less than Rascón, Rancho Viejo, and Micos surface fish (Fig. 2A). Clustering of surface, hybrid, and cave groups revealed that hybrid and cave populations slept significantly less than surface fish during the day (Fig. 2B,C). Taken together, these findings reveal a widespread evolutionary convergence on sleep loss in wild-caught populations of *A. mexican*us.

**Figure 2.**
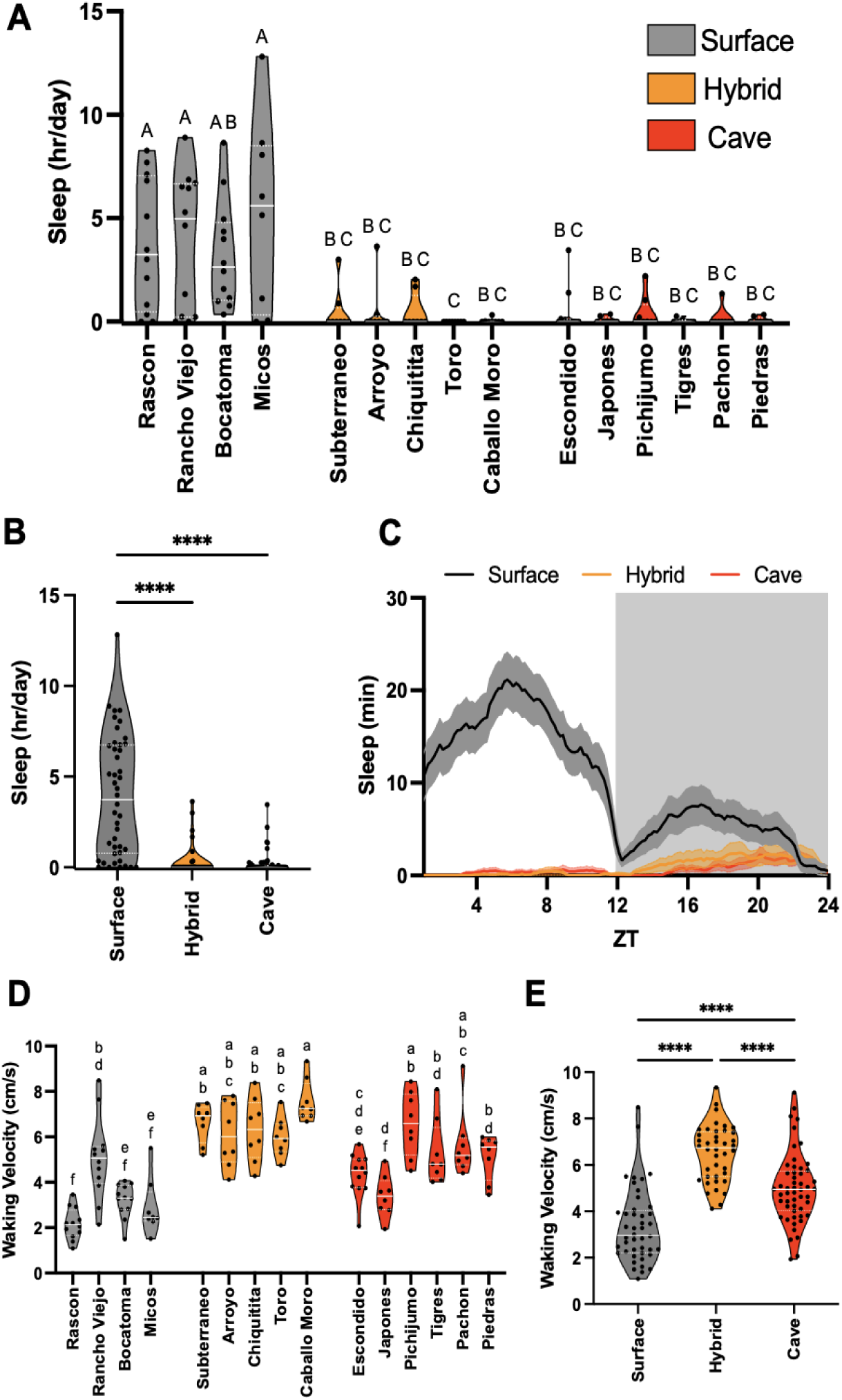
A) Total sleep (hr/day) across each independent population of Astyanax mexicanus. Sleep in Rascon, Rancho Viejo, and Micos was greater than all hybrid and cave populations (p < 0.001). **B)** Total sleep (hr/day) of all independent surface, hybrid, and cave populations grouped into their respective categories. When grouped, surface populations slept significantly more than both hybrid (p < 0.0001) and cave (p < 0.0001) populations, while cave and hybrid groups slept similarly (p = 0.9810). **C)** Sleep profile (mins) over a 24-hour period in a 12:12 light-dark cycle. Sleep duration over time from all individuals of surface, hybrid, and cave populations were averaged together. **D)** Average waking activity (cm/s) across each independent population. Waking activity in Rascon, Bocatoma, and Micos was lower than all hybrid populations (p < 0.001) as well as all cave populations except for Japones (p < 0.05; p < 0.005). **E)** Grouped population average waking activity (cm/s) of surface, hybrid, and cave categories. Grouped hybrid populations had significantly higher waking velocity than grouped surface (p < 0.0001) and cave (p < 0.0001) populations. Grouped cave populations had significantly higher waking velocity than grouped surface populations (p < 0.0001).

Reduced sleep is hypothesized to have adaptive value by allowing for increased foraging time in a nutrient-poor environment (Duboué and Keene, 2015; Horne, 2009). Indeed, previous studies in lab-reared fish revealed significantly greater locomotion and foraging-associated behaviors in cavefish (Duboué et al., 2011; Espinasa et al., 2014; Lloyd et al., 2018; Oliva et al., 2022). To examine whether increased activity is present across cave and hybrid populations of *A. mexicanus*, we quantified waking activity by normalizing the total activity to the wake duration (Duboué et al., 2011). The average waking activity for Rascon, Bocatoma, and Micos surface populations was lower than the waking activity of all hybrid populations, whereas the waking activity of Rancho Viejo surface fish was significantly lower than the Caballo Moro hybrid population (Fig 2D). Furthermore, the waking activity of all hybrid populations was greater than Escondido and Japones cave populations (Fig 2D). To further assess differences across surface, hybrid, and cave populations of *A. mexicanus*, we examined the combined phenotypes for each group. Waking activity was significantly greater in cavefish, compared to surface fish, while hybrid fish had significantly higher waking activity than both cave and surface fish (Fig 2E). Together, these findings reveal heterosis, or enhanced phenotypes in hybrids, for waking activity across lineages of *A. mexicanus*, and reveal differences in waking activity and sleep between hybrid and pure cave populations.

### Mapping behavioral and phenotypic traits on the phylogeny

Mapping complex traits such as sleep and waking activity can help us track their evolution and their association with ecological variation The presence of sleep loss and elevated waking activity provides the opportunity to examine the relationship between these behavioral traits and morphological traits commonly associated with cave evolution. We mapped sleep loss and waking velocity onto a phylogenetic tree based on the most recent phylogenetic reconstructions (Fig. S1A). This mapping revealed a clear pattern of repeated evolution in sleep loss and increased waking activity for all cave and hybrid populations, including at least three independent origins for sleep loss and elevated activity (Fig. S1A). In this regard, we observed that waking velocity was more variable than sleep loss across the cave and hybrid populations (Fig. S1A). Río Subterraneo cave showed a near complete loss of sleep with waking velocity amongst the lowest values, even lower than surface populations. For Japonés, in Sierra de El Abra, we observed the complete loss of sleep, but it was not accompanied by an increase in waking activity. When grouped by ecotype, the hybrid populations showed the highest waking velocity (Fig. 3B), while sleep was constant across cave and hybrid populations. Therefore, these findings confirm that the repeated evolution of sleep loss across *A. mexicanus* cave population, arose from evolutionarily distinct surface fish stocks.

**Figure 3.**
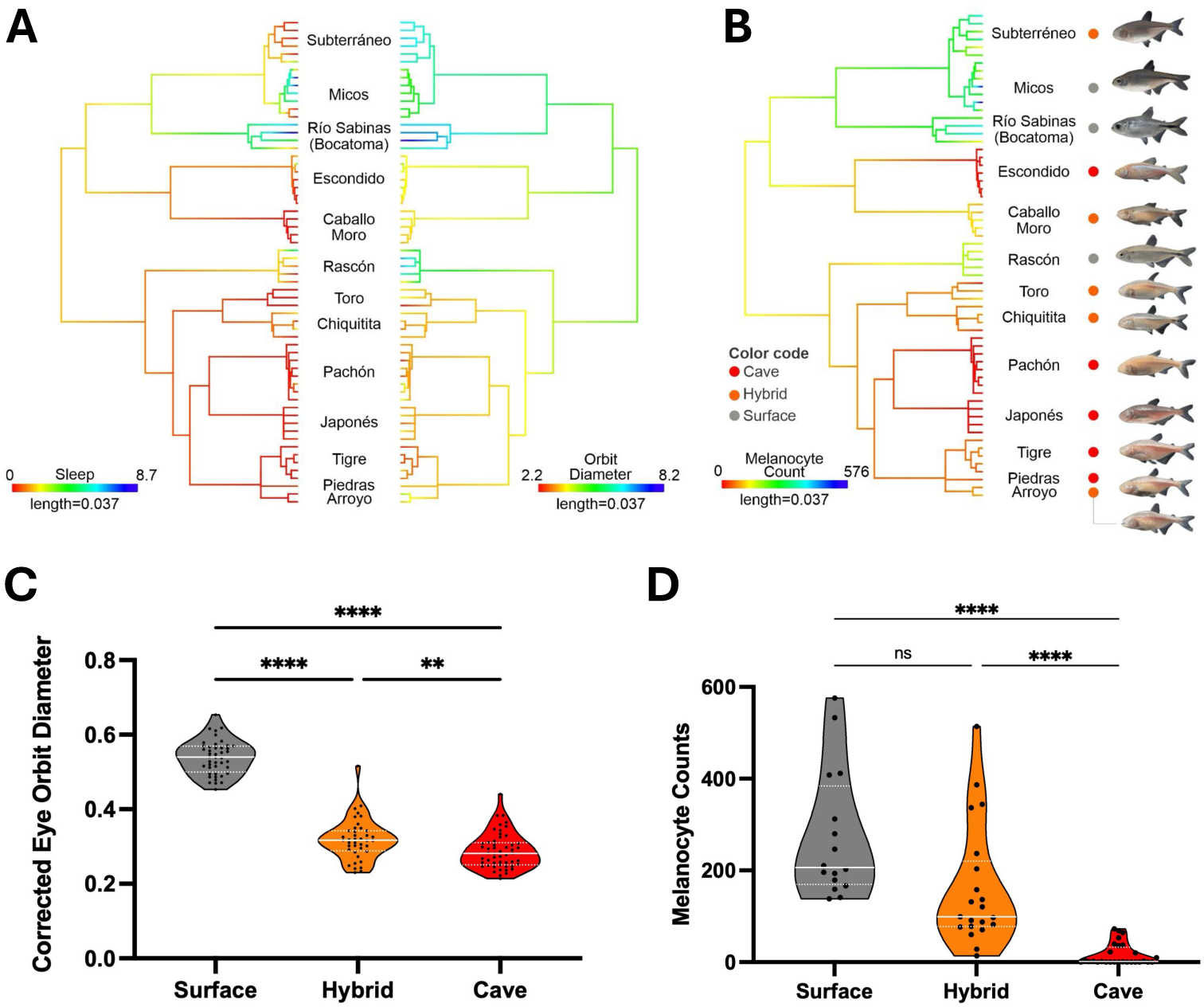
A) Sleep and Orbit Diameter mapped on a maximum likelihood phylogenetic tree based on 150,260 SNPs for 62 terminals, corresponding to the populations used in the study. **B)** Melanocyte count mapped on the ML phylogenetic tree. **C)** Average eye height adjusted by pre-opercular length values for grouped populations. Grouped surface fish had significantly larger corrected eye sizes compared to both hybrid (p < 0.0001) and cave (p < 0.0001) ecotypes. Grouped hybrid fish had significantly larger corrected eye sizes than grouped cave (p = 0.0082) ecotypes. **D)** Melanocyte count standardized by the dorsal fin length (melanocytes/mm). Grouped surface and hybrid fish had significantly more melanocytes than grouped cave (p < 0.0001) ecotypes.

We next sought to compare changes in sleep to morphological adaptations that are common in cave populations. Hybrid populations were intermediate in eye diameter size, with significantly more variable and larger eyes than non-admixed cave populations, and significantly smaller eyes than surface populations (Fig. S1B). Across hybrid populations, sleep loss was apparent in cases with only moderate reductions in eye size (Fig. 3A,C). When contrasting the melanocyte counts with sleep loss, hybrid populations showed the presence of pigment, but displayed extreme reductions in total sleep (i.e., Río Subterraneo, Caballo Moro, and Arroyo caves) (Fig 3B,D). Therefore, while sleep has been nearly completely lost in hybrid and cave populations, morphological phenotypes remain variable in regressive traits such as pigment and orbit diameter. Together, these findings reveal the near-complete loss of sleep and elevated waking activity in hybrid fish that retain, to a greater degree, surface-fish-like morphological traits, suggesting sleep loss and increased waking activity may be particularly potent drivers of cave evolution.

To date, the evolution of cavefish behavior has been studied extensively in the laboratory, but little is known about the behavior of these fish under natural conditions (McGaugh et al., 2020). While short-term behavioral measurements of sensory responsiveness have been performed in a cave environment, studies have not examined behavior for longer periods of time in cave or surface fish (Bibliowicz et al., 2013). To validate the ecological relevance of lab-based findings, we confined wild Rascón surface fish or Pachón cavefish to individual netting enclosures and recorded activity over 24 hours in their natural habitats (Fig 4A,B). Imaging limitations including debris and sediment accumulation prohibited the use of automated analysis systems that are regularly used for laboratory studies. Moreover, unlike the laboratory setting, fish movement was often passive due to currents, preventing the use of absolute activity as a characteristic of wakefulness. Therefore, we defined periods of active swimming, defined by movement of the tail fin, inactivity, defined by lack of active movement for 60 seconds or longer, and periods where the fish was obscured, defined as when the fish was not observed for 60 seconds or longer. These behaviors were manually scored using Behavioral Observation Research Interactive Software (BORIS) for a 24-hour period (Friard and Gamba, 2016), and sleep was defined as periods of inactivity 60s or longer. Similar to results from lab-raised fish, sleep is significantly reduced in cavefish compared to surface fish (Fig 4C-D). Analysis of sleep profiles revealed in the wild- setting, Rascón surface fish sleep more during the day than night, with no observable sleep in Pachón cavefish. These findings provide the first confirmation of sleep loss in any wild fish species and validate laboratory recordings from lab-reared and wild-caught fish.

**Figure 4.**
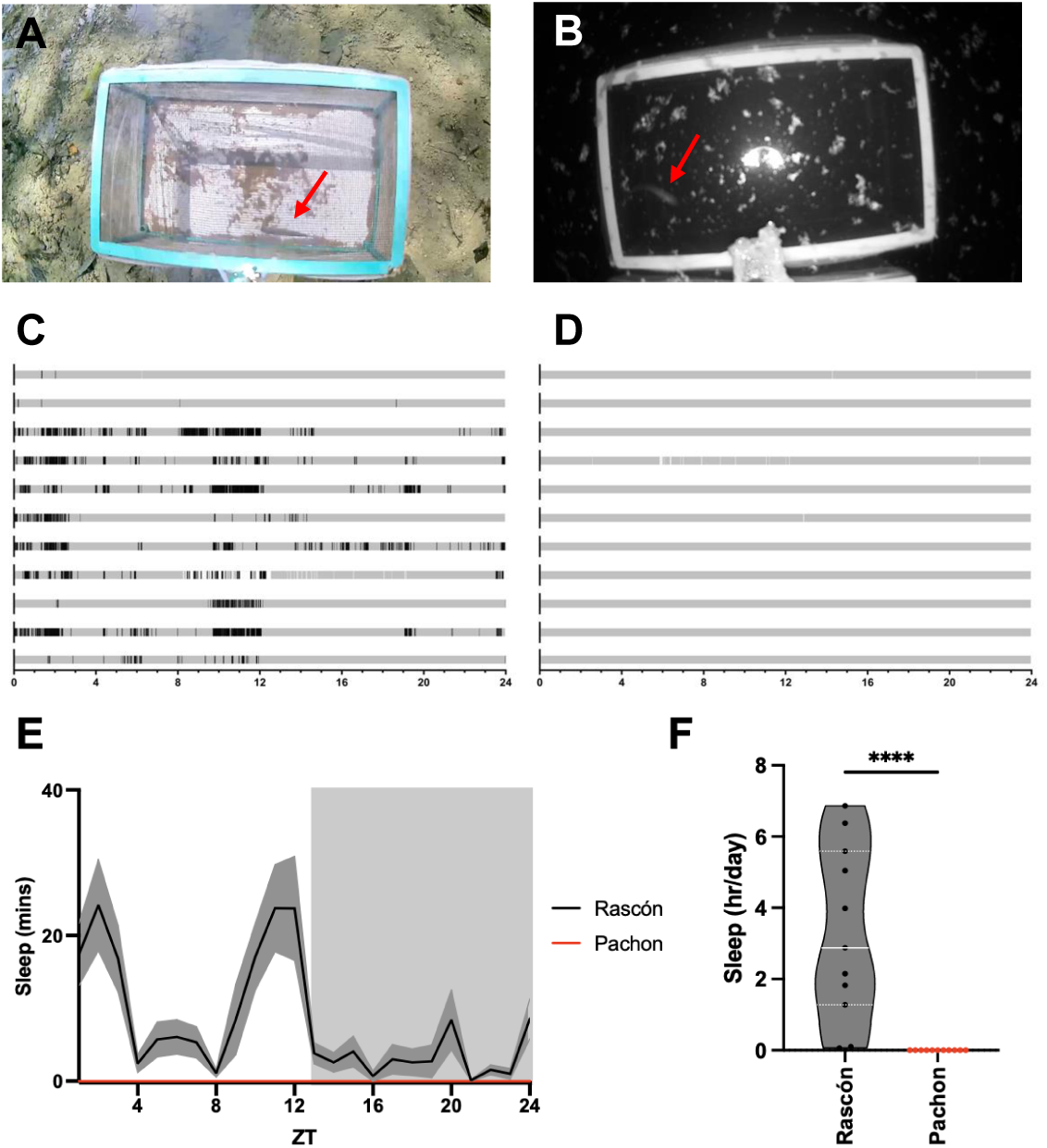
A,B) Images depicting a wild fish sleep recording setup from the Río Frio near Rascón, Mexico (A) and Pachón cave (B). Red arrows point to the single-housed fish in each arena. **C,D)** Manually generated ethograms of Rascón surface fish behavior (C)and Pachón cavefish behavior (D) Black areas denote periods of activity, grey denotes sleep, white areas denote periods where fish was obscured. **E)** Sleep profile over 24 hours captured in the wild reveals reduced sleep in Pachón cavefish compared to Rascón surface fish. **F)** Average total sleep (hrs/day) of wild fish from the Río Frio near Rascón, Mexico and Pachón cave. Wild Rascón individuals slept significantly more than wild Pachón cave individuals (p < 0.0001).

## Discussion

Our findings reveal repeated evolution of sleep loss in *A. mexicanus* cavefish lineages. Several species have been identified that forego sleep under specific contexts. For example, pectoral sandpipers suppress sleep seasonally during the mating season, while cetacean calves and their mothers sleep significantly less after birth (Lesku et al., 2012; Lyamin et al., 2008; Lyamin et al., 2018). Other animals including fur seals and frigate birds have developed unihemispheric sleep that allows for continuous activity during times when sleep may be unfavorable (Aulsebrook et al., 2016; Rattenborg et al., 2016). While these findings suggest numerous species possess the capability to forgo sleep, *A. mexicanus* are unique since sleep loss occurs across both development and context and has significantly diverged between two ecotypes (Duboué et al., 2011; Yoshizawa et al., 2015).

Here, we quantified sleep as 60 seconds of immobility, based on a previous study in adults that used this measure because it was associated with elevated arousal threshold and species-specific posture (Yoshizawa et al., 2015). Defining arousal threshold across *A. mexicanus* populations is confounded by differences in visual, olfactory, and mechanosensory processing.

For this reason, in a previous study we chose to use the response to mild electric shock, which elicits a robust behavioral response in surface fish (Yoshizawa et al., 2015). Similarly, mechanical tapping was used to define sleep in both surface fish and Pachón cavefish (Duboué et al., 2011). These methods suggest that similar periods of immobility are associated with sleep development and using multiple sensory stimuli, supporting the notion that the period of immobility used to define sleep does not need to be established for each population.

While the application of this behavioral analysis allows for efficient scoring of sleep, it is possible that the time of immobility indicative of sleep differs across populations, and this analysis does not rule out confounds associated with differences in sensory processing. In larval zebrafish, genetically encoded Ca^2+^ sensors have been used to define sleep, providing precise measures of REM-like and non-REM-like sleep (Leung et al., 2019). We have developed these transgenic cavefish harboring these sensors, and therefore it is feasible, though challenging, to examine physiological readouts of sleep (Lloyd et al., 2021). These studies would be more challenging in adults that lack a transparent skull, though studies in adult zebrafish have applied Ca^2+^ imaging, or implanted electrodes. Therefore, application of additional methodology to study sleep may further inform the biology underlying the repeated evolution of sleep loss.

A finding from this analysis is that there is increased waking activity in hybrid cavefish, compared to non-admixed cave populations. This raises the possibility that elevated locomotion provides a competitive advantage to hybrids through enhancement of foraging. Examples of heterosis, or hybrid vigor, are relatively rare in the wild, and have not been reported for sleep or locomotor activity (Bar-Zvi et al., 2017). Sleep and waking activity are highly pleiotropic, suggesting that interactions between complex gene regulatory networks contribute to heterosis. Previous work has speculated that sleep loss and increased activity represent adaptations to improve foraging in nutrient-poor cave environments (Duboué and Keene, 2015; Duboué et al., 2011; Keene and Duboue, 2018). It is possible that the waking activity in hybrids represents an intermediate step that allows for improved foraging, prior to the evolution of secondary traits that improve survival in caves. For example, a suite of cave traits support energy conservation and survival during periods of prolonged food deprivation (Krishnan et al., 2022; Pozo-Morales et al., 2024; Volkoff, 2016), and may reduce the selective pressure for such significant increases in waking activity.

Phylogenetic comparative methods allow us to evaluate the patterns of trait evolution (Revel, 2023), such as sleep and vision loss. Through the mapping of a trait’s evolutionary history, we can make inferences about the timing and order of trait evolution, particularly in species with defined populations that diverged (Schluter et al., 1997). Moreover, there are trade-offs between behavior and morphological variation that could affect their macroevolutionary patterns. This study is the first to examine behavioral evolution across approximately a third of cavefish populations, and map changes onto a known phylogeny. Based on our mapping, sleep loss and increased waking activity have evolved independently among the different cave lineages. Although vision loss and pigmentation have also evolved in parallel among cave populations, these traits show greater variation than sleep and waking activity, particularly in hybrid populations. We used a random walk evolutionary model, which represents an unpredictable evolutionary scenario not directed by natural selection; this method does not imply that evolution is random (Schluter et al., 1997). Alternative models, such as Bayesian ancestral character estimation or GLS-OU, were not tested due to sampling limitations, which introduce substantial uncertainty into ancestral trait estimation(Revell, 2024). However, our results showed consistently a convergent evolutionary pattern of sleep loss and waking velocity in cave populations, whose lineages are well represented in this study.

Prior to this study, all analyses of sleep in *A. mexicanus* compared sleep in cave populations to a single surface population, Rio Choy. Testing four additional surface populations provided the opportunity to map changes in sleep across the well-defined phylogeny of *A. mexicanus.* This analysis revealed that sleep loss and elevated waking activity have occurred at least three times over the course of evolution. These findings raise the possibility that evolutionarily distinct mechanisms contribute to sleep loss across populations. To date, there is some lab-based experimental evidence to support this notion. For example, the *oculocutaneous albinism type 2* (*oca2)* is the genetic cause of albinism across multiple cavefish populations, and mutation of *oca2* in Rio Choy leads to sleep loss (O’Gorman et al., 2021). However, there is also robust sleep loss in Tinaja cavefish that maintain pigmentation and *oca2* function. Similarly, ablation of the lateral line in Pachón cavefish restores sleep to surface fish levels, but does not change sleep in Molino, Tinaja, or Chica cavefish (Jaggard et al., 2017). These findings suggest increased sensory input from the lateral is required for sleep loss in some, but not all, populations of *A. mexicanus* (Jaggard et al., 2017). Defining multiple lineages with sleep loss provides the opportunity for future mechanistic studies and complementation analysis examining whether shared, or distinct processes contribute to sleep loss.

The identification of such widespread loss of sleep among cavefish populations has potential to offer insight into the processes that shape evolution. Food availability is thought to be limited compared to the surface environment, which could increase selective pressure for improved foraging. Another proposed function of sleep is predator avoidance. Cave populations of *A. mexicanus* have no reported predators, raising the possibility that reduced selective pressure may contribute to the loss of sleep. Nevertheless, sleep is thought to play a fundamental role in the maintenance of physiological homeostasis and immune function, and prolonged sleep loss leads to death in fruit flies and mammals (Rechtschaffen and Bergmann, 1995; Vaccaro et al., 2020). Laboratory studies have shown that cavefish harbor markers of DNA damage and reactive oxygen species, both indicators of chronic sleep loss. Therefore, the ecological factors contributing to the evolution of sleep loss, and their biological consequences, remain to be determined.

Taken together, these findings reveal sleep loss is broadly conserved across cave and hybrid populations. This work also represents the first study to compare sleep in wild-caught fish from cave populations to fish from multiple independent populations of surface fish, allowing for the evolutionary origins of sleep loss to be determined. Our findings demonstrate that sleep loss has arisen at least three times across Mexican cavefish evolution. To our knowledge, *A. mexicanus* cavefish remains the only species identified to date with nearly complete sleep loss across its life cycle (Keene and Duboue, 2018). Since these populations of cave morphs arose from cave invasions by independent surface stocks, and have evolved repeatedly over different periods of time, it is likely that sleep loss is critical for fitness in the cave system. The identification of sleep loss in these populations provides the opportunity to test the physiological and functional trade- offs between sleeping and non-sleeping populations of *A. mexicanus*, with potential to provide insights into the long-elusive evolutionary function of sleep.

## Supporting information

Supplemental Statistics

Supplemental animal information

## Acknowledgements

This work was supported by the National Institute of Health R24OB030214 to ACK and R21 NS122166 to ACK and US-Israel Binational Science Foundation Award 2021177 to ACK and NSF IOS 2202359/1933076 to JEK and SEM. This research was also supported by the PAPIIT UNAM No. IN214624 to CPOG.

## Supplemental Methods

### Husbandry and Video Recordings

We included *A. mexicanus* specimens from a total of 15 different localities from the Sierra de El Abra (i.e, Pachón, Japonés, Piedras, Tigre, Arroyo, Pichijumo, Chiquitita and Toro), Sierra de Guatemala (i.e., Caballo Moro and Escondido), and Sierra de La Colmena (i.e, Micos) regions, and four surface populations: Bocatoma, Rancho Viejo, Subterráneo and Rascón from regions in the states of San Luis Potosí and Tamaulipas in Mexico (Fig 1B). These specimens were collected during various field trips conducted from 2015 to 2020. Collection permits were obtained from the relevant authorities (SEMARNAT SGPA/DGVS/2438/15-16, SGPA/DGVS/05389/17, and SGPA/DGVS/1893/19). The fish were kept alive and maintained in our facility at the Institute of Biology, UNAM (CNPE, IB-UNAM, Mexico City). All animals were housed at the fish facility at 22°C ± 1°C. Two weeks prior to the experiments, animals were fed a mixture of dry and live food diets (i.e. commercial flakes and blood worms). The protocol was approved by the Comisión de ética académica y responsabilidad científica, Science Faculty, UNAM.

For behavioral assays, we established a tracking system based on previously described methods(Moran et al., 2022b). Individual fish were placed in 20 L tanks fitted with custom-cut white corrugated plastic (3 mm thick) illuminated with constant infrared light, and 12:12 LD white light. Following a 24 hrs acclimation period, fish were recorded for a period of 24 hrs using a Microsoft LifeCam Studio 1080p HD Webcam - Gray, fitted with an IR lens (Quanmin 9.6mm×1.0 mm Slim Optical 780nm Filter Infrared Cold Mirror). Locomotor behavior was tracked in Ethovision XT 17.0 (Noldus), and frame-by-frame velocity data was exported and analyzed using a custom MATLAB script, with sleep defined as 60 seconds or more of consolidated immobility. A velocity cut-off of 4 cm/s was used to distinguish active swimming from passive drift. Data was processed using custom-written MATLAB scripts as previously described (Moran et al., 2022b).

### Morphological Phenotyping

After the behavior recordings, fish were anesthetized in cold water for 30 sec. The fish were photographed from the left side with a reference scale. Subsequently, using a stereomicroscope (Stemi 305, ZEISS ZEN), a photograph was taken of the left side of the head of the specimen with a size scale. Using ImageJ v.1.54r (Schindelin et al., 2012) the following measurements were taken: 1) orbit diameter (mm), and 2) head length (mm) from the tip of the mouth to the posterior edge of the pre-operculum. The orbit diameter dimensions were corrected using the head length (mm). A variance analysis was performed to compare the mean differences between the ecotypes (i.e., caves, surface, and hybrids). Kruskal-Wallis and a pairwise Wilcoxon rank test were used to compare orbit diameter across ecotypes.

### Pigmentation measurements

Photographs of the left side of the fish were obtained using a stereomicroscope (Stemi 305, ZEISS ZEN), with a size scale. Using ImageJ v.1.54r (Schindelin et al., 2012), a line was drawn at the base of the dorsal fin, with the Straight-Line Tool, to define the fin length. The value obtained was used as a reference for the definition of a grid, dividing the length of the fin base into 7 squares. Melanocytes were quantified in four of the seven squares in the first line below the dorsal fin, and the mean for the four squares per individual was obtained. Two selected squares corresponded to the extremes of the dorsal fin base, and the remaining two, to the central area of the fin base. The same scheme was repeated for all the analyzed specimens.

To improve the detection of melanocytes, the images were preprocessed with the Sharpen tool, which improved the contrast of the images. Subsequently, with the Find Maxima function, the points of maximum intensity in the image were identified. A range between 10 and 15 pixel intensity units was set to ensure correct identification of the melanocytes in the area, without including illumination artifacts. Once the threshold was adjusted, an automatic count of melanocytes in the selected area was obtained. To visualize the total number of cells identified and to refine the segmentation, the Maxima Within Tolerance option within Find Maxima was applied. Finally, a quantitative particle analysis was performed using the Analyze Particles function. This analysis generated a summary of the total number of melanocytes detected.

### Mapping behavioral and phenotypic traits on the phylogeny

A phylogenetic reconstruction was performed based on RADseq data from wild-caught and previously published complete genomes (Garduño-Sánchez et al., 2023; Moran et al., 2023). The samples used as a reference for the Bocatoma population corresponded to the Rio Sabinas population (Garduño-Sánchez et al., 2023), because they come from the same river basin. The bioinformatic processing methodology followed protocols similar to those previously described (Garduño-Sánchez et al., 2023). In total, 188 samples were downloaded from the Sequence Read Archive (SRA), for the species *Astyanax mexicanus*, two specimens of *A. aeneus* and two of *A. nicaraguensis*, collected in Nicaragua as an outgroup (Garduño-Sánchez et al., 2023; Herman et al., 2018). The quality of the reads was assessed using FastQC, and low-quality reads, as well as adapters, were removed using Trimmomatic (Bolger et al., 2014). Subsequently, the processed reads were aligned against the most recent reference genome of *A. mexicanus* available at NCBI (GCF_023375975.1) using the BWA-MEM2 aligner (Vasimuddin et al., 2019). Only Rancho Viejo and Pichijumo were not included in the phylogenetic reconstruction due to the lack of genetic information.

Genotyping and variant calling were performed using BCFtools.(Danecek et al., 2021). Quality filters were applied with VCFtools (Danecek et al., 2021), allowing up to 30% missing data, excluding indels, and retaining SNPs with a minimum mapping quality of 50 and a minimum depth of 7 reads. In addition, alleles with a minimum frequency of less than 0.01 were removed. After bioinformatic processing of the NGS samples, the final dataset consists of a total of 150,260 SNPs. These markers were obtained from 172 samples of *Astyanax mexicanus*, from cave and surface populations, as well as from two samples of *A. aeneus* and two specimens from Nicaragua, used as outgroups.

With the SNP’s obtained after filtering, phylogenetic reconstruction was carried out through IQ-TREE (Minh et al., 2020). The VCF format file with the 192 total samples was transformed to FASTA format using the vcf2phylip program. For phylogenetic inference, an initial search was performed on 1000 parsimony trees by random stepwise addition and pruning of subtrees (Flouri et al., 2015). The 10 best-resulting trees were used for maximum likelihood-based reconstruction by applying the GTR+G substitution model and the ascertainment bias correction model (+ASC; Lewis, 2001) for data without constant sites. Branch support was assessed using ultrafast bootstrap (UFBoot;(Hoang et al., 2018)) and the SH-aLRT test (Guindon et al., 2010), each with 10,000 replicates. To minimize the overestimation of branch supports due to model violations, the nearest neighbor exchange (NNI) search was optimized for each bootstrap replicate using the - bnni option in IQ-TREE.

To conduct the trait mapping, the phylogeny previously calculated was pruned to the populations that were not included in our behavioral study using ape version 5.7.-1. The final topology included 62 individuals corresponding to 13 populations analyzed. To visualize the evolutionary patterns, we mapped the following data on the phylogeny: 1) raw sleep data, 2) waking velocity, 3) corrected orbit diameter by head length, and 4) melanocytes counts. Estimation of node internal values was conducted using the fastAnc and trait visualizations using the contMap function, both included in phytools version 2.1-1in R-Studio v. 4.02.

### Analysis of fish in the natural environment

Surface fish were assayed from the Río Frio near Rascón, Mexico and cavefish were assayed from Pachón cave in Mexico between March 18 - March 26 2022. Fish were caught using collapsible bait traps and sardines as bait. Each fish was transferred to its individual breeder box (Nylon Mesh Fish Fry Hatchery Breeder Box Separation; 26.5 x 15 x 15.5cm, L x W x H) which was modified to have a white plastic poster board bottom to better visualize the fish, and which had the top replaced with screen door mesh to avoid fish escapes. Breeder boxes were mounted to garden stakes and submerged so the top of the breeder box was just below the water surface. Outdoor Wyze Cam v1 cameras were mounted above the breeder boxes, using one camera per fish/arena and visualized to ensure the breeder box was encompassed within the camera frame. Outdoor Wyze Cams were set to auto mode to turn on infrared and infrared filters to record in night vision mode once the camera detected sundown. Cameras continuously recorded in 1080p resolution to a 32Gb microSD card. Renogy 72000mAh Laptop Power Banks were used to power the cameras beyond the camera’s battery life. The power banks required a small load to be pulled to stay in charging mode until the cameras’ batteries were exhausted and then the camera started to pull the load. A FINENIC 6-Black Mini USB LED Light Lamp wrapped in aluminum tape to completely block out any light coming from the lamp was plugged into a VENTION 4-Port USB 3.0 Hub Ultra-Slim Data USB Splitter to pull this small load and keep the Power Bank from turning off. After recording concluded, fish were photographed with a ruler in frame. Methods were approved by University of Minnesota.

We analyzed the behavior of fish for a continuous 24-hour period beginning at 6:40am local time, coinciding with the return of day mode recording settings, and ending at 6:40am the following day. The beginning of the analysis period corresponds to local sunrise, which occurred f 6:35am - 6:41am, depending on location and date of recording. Fish were acclimated within the arena for at least 14 hours before analysis began. Videos were annotated for when the fish was performing non-active swimming and when the fish was obscured using the open-source software BORIS (v.8.27.1) (Friard and Gamba, 2016)

Non-active swimming was scored as the fish not actively making body movements for a period longer than 20 seconds. This metric included periods of drift due to river flow during which the fish slowly drifted but did not actively engage in swimming movements. Bouts of non-active swimming conclude with a sharp, distinct change in orientation or a sudden increase in velocity. Small adjustments to the fish’s positioning did not end a bout of non-active swimming. Due to the unique challenges of filming in the field, fish were difficult to observe for portions of the 24-hour period. Fish behavior was scored as obscured when the fish was not sufficiently visible to make an accurate assessment of the positioning of the fish for a period of 20 seconds or longer. For example, if a fish was observed in active swimming at t=1 sec and t=19 sec, but obscured in between, the fish was scored as active swimming rather than obscured.

The behavior of 11 Rascón surface fish and 11 Pachón cavefish was analyzed for 24 hours. Following analysis, observation time stamps were exported to a .csv in BORIS. Even though behaviors were annotated after 20 seconds, only observations of inactivity longer than 60 seconds were considered for further analysis.

### Statistics

All statistical tests of behavior were conducted using InStat software (GraphPad Prism 10), with p < 0.05 set as the threshold for significance. An ordinary one-way ANOVA test followed by a Tukey’s multiple comparison was used to test for significant differences in sleep between the 15 independent populations of wild-caught surface, hybrid, and cave fish (Fig 2A). To test for differences in total sleep between grouped surface, hybrid, and cave populations, an ordinary one-way ANOVA test followed by a Tukey’s multiple comparison was conducted (Fig 2B). To test for differences in waking velocity across independent populations, an ordinary one-way ANOVA test followed by a Tukey’s multiple comparison was conducted (Fig 2D). To test for differences between waking velocity in grouped surface, hybrid, and cave populations, an ordinary one-way ANOVA test followed by a Tukey’s multiple comparison was conducted (Fig 2E). To analyze differences and variability in eye size, the average eye orbit diameter of each individual (mm) was standardized by the length of their head (mm), defined as “corrected eye orbit diameter”. To test for significant differences in corrected eye orbit diameter, an ordinary one-way ANOVA was performed, followed by Tukey’s multiple comparison (Fig 3C). To analyze differences in melanocyte counts across populations, melanocytes were quantified and standardized by dorsal fin length (melanocytes/mm), defined as “melanocyte counts”. To test for significant differences in melanocyte counts, a Kruskal-Wallis test was conducted, followed by a Tukey’s multiple comparison (Fig 3D). To analyze significant differences in sleep duration between wild individuals recorded in the Río Frio near Rascón, Mexico, and the Pachon cave, a Mann-Whitney T-test was used (Fig 4F).

---

**Supplemental Figure 1.**
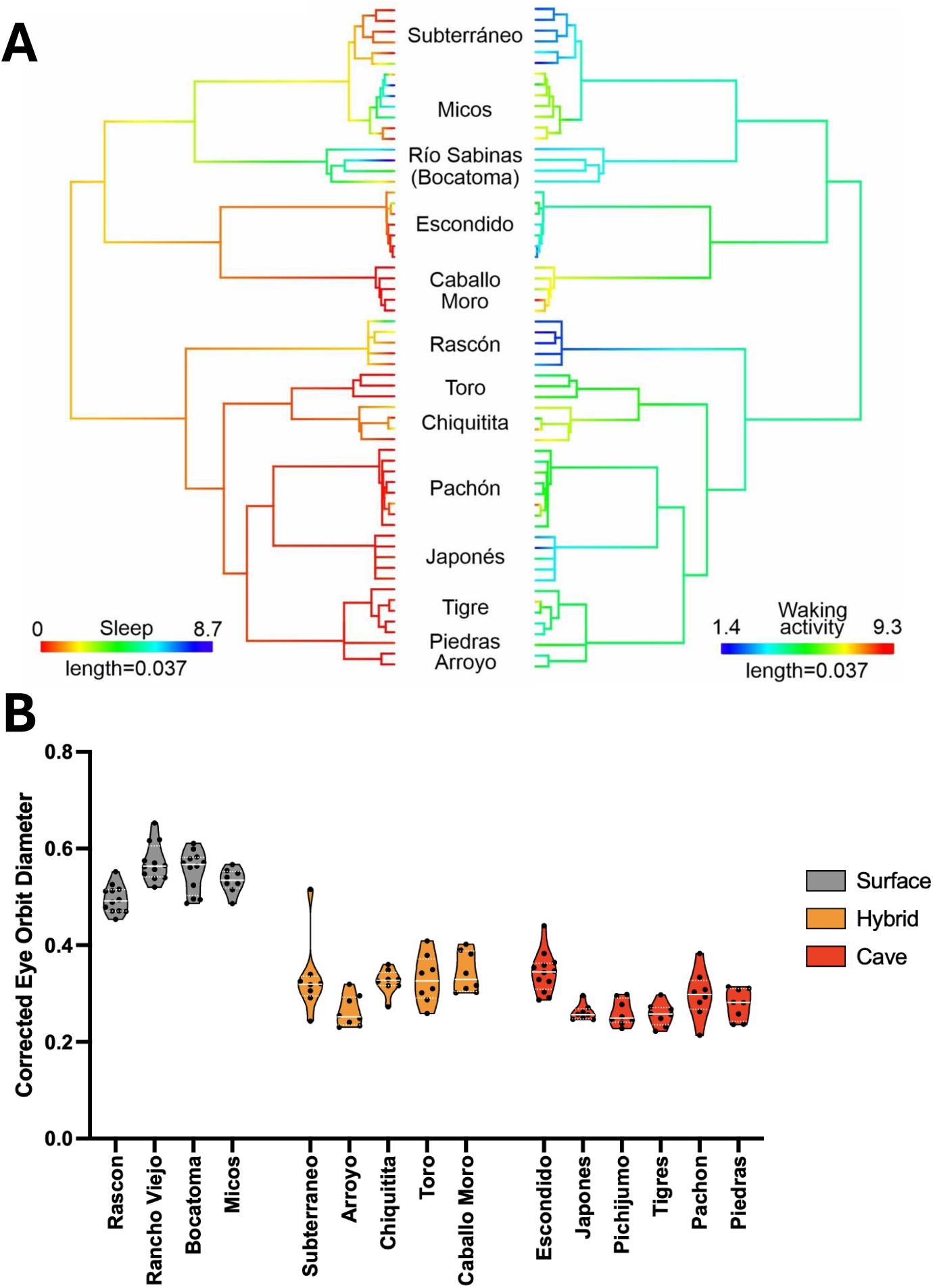
A) Sleep and waking activity mapped on a maximum likelihood phylogenetic tree based on 150,260 SNPs for 62 terminals, corresponding to the populations used in the study. **B)** Corrected eye orbit diameter across each independent population tested in the study.

